# Feline tooth resorption in a case-control study based on a subpopulation of 944 dentally examined cats from a Finnish questionnaire survey of over 8000 cats

**DOI:** 10.1101/2021.01.22.427753

**Authors:** Katariina Vapalahti, Henriikka Neittaanmäki, Hannes Lohi, Anna-Maija Virtala

**Affiliations:** Department of Veterinary Biosciences, 00014 University of Helsinki, Finland; Department of Medical and Clinical Genetics, 00014 University of Helsinki, Finland; Folkhälsan Research Center, Haartmaninkatu 8, 00290, Helsinki, Finland

**Keywords:** cat, feline, tooth resorption, FORL, dental, oral disease

## Abstract

Tooth resorption (TR) is one of cats’ most common dental diseases. It is a painful condition characterized by progressive dental destruction, which eventually results in loss of teeth. The aetiology of the TR remains unclear, but associations with old age, breed, other oral and dental diseases, and certain environmental factors have been suspected. In our study, we used part of the data from the extensive feline health and environmental survey of 8115 Finnish cats collected through an online survey targeted at cat owners. We aimed to investigate the characteristics of cats having TR and to study risk factors for TR. Because TR is difficult to detect and, in addition, the feline health survey was very comprehensive and included diagnoses defined by both veterinarians and owners themselves, we limited our study to a subpopulation of cats diagnosed with oral or dental disease by a veterinarian and undergone dental examination or surgery under sedation (n=944).

The frequency of veterinary-diagnosed TR was 3.9% in the entire health survey data (316/8115) and increased to 21% in the subpopulation of veterinarian-diagnosed and sedated cats (202/944). We utilized case-control multivariable logistic regression in this subpopulation to determine the risk factors and breed variation of feline TR. The 202 cats diagnosed with TR were defined as TR cases and the 742 cats without TR diagnoses served as controls. Results indicate that the risk of TR increases with age. Dental calculus, gingivitis, and periodontitis were associated with TR. These findings and the interaction of dental calculus with gingivitis and periodontitis might suggest that inflammatory changes caused by dental calculus increase the risk of TR. We found Cornish Rex, European, and Ragdoll at higher risk for TR. Exotic-Persians had lower risk, and Turkish van and Devon Rex had no TR. The observed differences between breeds highlight a genetic contribution. In addition, female cats that had food available constantly had significantly less TR than female cats that had feeding times. The underlying influential reasons for this result remain unexplained in our study.

## Introduction

Tooth resorption (TR) is one of cats’ most common dental diseases. It is a painful condition characterized by progressive dental destruction (Palmeira et al., 2022), which eventually results in loss of teeth. Dental radiography is necessary to evaluate the overall situation (Reiter and Mendoza, 2002). Prevention of the disease is not possible since the aetiology is still unknown. The goal of treatment is to relieve the pain and discomfort caused by these lesions (Gorrel, 2008). The leading cause of destruction is odontoclasts, multinuclear cells that resorb mineralized tissue (Okuda and Harvey, 1992). Odontoclasts are responsible for the resorption of deciduous teeth in young animals, but their abnormal activity in permanent teeth is the cause of TR (Scarlett et al., 1999). The reason for this process remains unclear, although many theories have been proposed. Tooth resorption can include plaque accumulation, inflammation of the adjacent tissue and alveolar bone ankylosis (DeLaurier et al., 2009). Because the pathogenesis of tooth resorption has been unclear, many different terms have been used to describe the lesion, i.e., erosion, neck lesion, FORL (feline odontoclastic resorptive lesion). Nowadays, the most used term is TR.

Increasing age has widely been found to increase the risk of TR (Coles, 1990; Harvey, 1993; Lund et al., 1998; Ingham et al., 2001; Pettersson and Mannerfelt, 2003; Reiter et al., 2005b; DeLaurier et al., 2009; Mestrinho et al., 2013; Pistor et al., 2023). Scarlett et al. (1999) reported that cats with previous dental diseases (gingivitis, calculus, or periodontal disease) had four to five times higher odds for resorptive lesions than those without the previous dental disease. Other studies have also found the association of gingivitis with TR lesions (Harvey, 1993; Mestrinho et al., 2013; Pistor et al., 2023). Some studies have not found an association with gingivitis (Ingham et al., 2001; Gorrel and Larsson, 2002). However, the methods of Gorrel and Larsson (2002) differed from others as they examined individual teeth histologically. Periodontitis has been associated with inflammatory tooth resorption (DuPont and DeBowes, 2002). However, tooth resorption can cause inflammatory resorption of the surrounding alveolar bone and increase the risk of decreased alveolar bone height (Lemmons, 2013). Farcas et al. (2014) did not find TR more in cats with chronic gingivostomatitis. The size and origin of the study population and methods have a major impact on the reported prevalence of TR in previous studies, varying from 29-85% (Coles, 1990; Lund et al., 1998; Petterson and Mannerfelt, 2003; DeLaurier et al., 2009; Whyte et al., 2020). Most studies are based on a small clinical sample (n< 150), and the studies with the biggest sample size are van Wessum (1992) with 432 cats and 62% prevalence, Lommer and Verstraete (2000), with 265 cats and 60.8% prevalence, and Ingham et al. (2001), with 228 cats and 29% prevalence.

Our aim was to study the risk factors for TR in a case-control study in a subset of a large cat health survey - cats with veterinary-diagnosed oral or dental disease and underwent dental examination or surgery under sedation. The subset was used to minimize the number of false positives and negatives in the data.

## Materials and Methods

The material for this study was part of the data that were obtained from a cross-sectional online feline health questionnaire survey targeted at all Finnish cat owners (Vapalahti et al., 2016). The questionnaire included the cats’ environment, diseases, and behaviour. The disorders were divided into various categories including several specific diseases. Each category also included information on whether the diagnosis was made by a veterinarian or by the owner. The questionnaire and data collection methods in detail and some results of the survey have been published previously by Vapalahti et al. (2016), Ahola et al. (2017), and Salonen et al. (2019).

Because TR is difficult to detect and the definitive diagnosis requires clinical examination and dental radiographs under sedation, we included in this study only those cats from the feline health survey that had undergone veterinary oral or dental examination or surgery under sedation (according to the open-ended field of question). For this subgroup, we defined those cats with a TR diagnosis as TR cats and the rest as non-TR cats. When evaluating breed associations, we used house cats as the reference group, as they represented the average cat population in terms of TR and other diseases (Vapalahti et al., 2016).

The data used in our study consisted of 1) basic information about the cat: breed, registration number, day of birth, possible day of death, sex, neutering, 2) environmental factors: vaccinations, outdoor habits, diet, home environment, and 3) disease categories and specific diagnoses. The specific diagnoses were considered only in oral and dental, and autoimmune diseases, and viral infections. The differential diagnoses in dental and oral diseases were malocclusion, gingivitis, stomatitis, periodontitis, tooth resorption (odontoclastic resorptive lesion FORL in Vapalahti et al. (2016)), dental calculus, tooth fracture, and an abnormal number of teeth.

The cat’s age was the difference between the date of birth and the response date. Age was categorized into three age groups: <7 years, 7 to < 11 years, and at least 11 years. We utilized breed groups of genetically similar breeds (presented in Vapalahti et al., 2016) in breeds with low numbers. The breed groups were Abyssinian–Ocicat–Somali, Siamese–Balinese–Oriental–Seychellois, Exotic–Persian, Cymric–Manx and Neva masquerade-Siberian. Only the veterinarian’s diagnoses were included in the study in the dental and oral disease category. In other disease categories, the initial options ‘veterinarian’s diagnosis’ or ‘own diagnosis’ were summed up to option ‘1’ (yes) if either option was selected. ‘Not known’ responses were coded as missing; the rest were ‘0’ (no) responses. Coding ‘1’ (yes) or ‘0’ (no) was used to analyse specific diagnoses. The environmental information, such as diet and vaccinations, were coded similarly. Outdoor habits were divided into five categories: ‘Outdoors every day’, ‘Outdoors all along’, ‘Outdoors weekly’, ‘Outdoors monthly’ and ‘Outdoors less often than monthly/never’. Genders were coded as ‘Female’ and ‘Male’. The effect of neutering was not studied since most cats were neutered.

We examined cats’ demographic and environmental factors separately in TR and non-TR cat groups. The proportion of tooth resorption in different breeds, age groups, sex, other diseases, and environmental factors were calculated with proc freq procedure (SAS 9.4, SAS Institute, Cary NC, 2016)). The binomial option was used to calculate Wilson 95% confidence intervals (Brown et al., 2001) for percentages. The statistical significance of factors associated with TR was evaluated with Fisher’s exact test for two categorical variables and Chi-square test for multicategorical variables by utilizing fisher or chisq option in proc freq procedure. Factors at level P <0.2 in the basic tests above were analyzed individually in preliminary logistic regression models (SAS proc logistic procedure) with confounding factors. Age, sex, and breed were considered confounding factors according to the literature (Houe et al., 2004). In the modelling, we treated TR-cats as cases and the rest of the cats as controls, and variable TR – ‘yes’ or ‘no’ – was set to a response variable with option ‘yes’ as the event. Variables with P-value <0.05 (Wald chi-square) in preliminary logistic regression models were qualified for the multivariable logistic regression modelling for TR’s most important risk factors.

The multivariable modelling (SAS proc logistic procedure) tested interactions until the second. The model selection was performed using the forward, backward and stepwise selections and choosing the best model of these according to goodness of fit statistics. P-value <0.05 of Wald chi-square was set to cut off value for covariate significance. The goodness of fit of the model was evaluated with Pearson goodness of fit statistics, McFadden index, Akaike information criterion (AIC), and the predictive value by the area under the curve (AUC) of the receiver operating characteristic (ROC) curve. Multicollinearity between variables was estimated by the Phi coefficient – the limit value for strong correlation was set at 0.5 (Pett, 1997).

As a comparative analysis with the logistic regression procedure, we performed random forest (RF) (Ho, 1995) classification using R (R Core Team, 2019) randomForest package (Liaw and Wiener, 2002). In order to make RF analysis data to be comparable with logistic regression analysis data in terms of observation numbers, we included in the RF data those variables that had adequate number (>4) feature occurrences in both TR-groups and less than 20 missing values. Eventually, the data included 31 variables. The importance of the variables in predicting TR was calculated in accuracy perspective. We set the number of trees to 500 in RF, and manually tuned the number of randomly drawn candidate variables and the minimum number of observations in a terminal node to find the best accuracy for the classification. The data division into to train data (80%) and test data (20%) and creation of importance plot of variables were performed using R caret package (Kuhn, 2022).

## Results

### Frequency of tooth resorption

The cat population of the original feline health survey (Vapalahti et al., 2016) consisted of 8115 cats in 40 breeds. Of the cats, 2070 (26%) were diagnosed with oral or dental disease, and 316 (3.9% (95% CI 3.5–4.3%)) cats had a veterinarian diagnosed with TR.

Restricting the cat population to cats that had veterinarian-diagnosed oral or dental disease, and which, according to the responder’s report (open ended-question), had undergone a veterinary oral or dental examination or surgery under sedation, reduced the cat population to 944 cats in 35 breeds; 202 cats with TR (21.4 %; 95% CI 18.9–24.1 %) and 742 without TR. Of the TR-cats, 169 had had teeth extractions done because of TR (84%), as clarified in the open-ended field of the question. The mean age of the cat population was 8.9 years – 8.6 years for cats without TR and 9.8 years for cats with TR. The frequency of TR increased by age, 14.7% in 0 to <7 years old, 25.9% in 7 to <11 years old, and 25.3% in at least 11 years old cats.

We found no significant difference between the TR frequency in purebred cats and non-pedigree house cats (21.5%; CI 18.8–24.6% and 20.8%; CI 15.7–27.1%, respectively, P = 0.922, Fisher). However, in some breeds, the frequency was much higher or lower than in the entire in-house cat population (21.4%) – high in Cornish Rex (45.1%), European (37.5), Ragdoll (35.1%) and in the Oriental group (31.9%), and much lower in Birmans (8.6%) and Russian blue (10.0%) and Persian-Exotic group (10.9%). TR was not reported in the breeds of Devon Rex and Turkish van.

### Factors associated with tooth resorption

We selected 51 factors, including the confounding factors age, sex, and breed, from the feline health survey (Vapalahti et al., 2016) that were previously shown to be or assumed possibly to be related to TR (Supplementary Table S1). In the basic association tests (Fisher’s exact and Chi-square), ten factors in addition to age, sex, and breed qualified for further analysis (P <0.2) (Table 1, Supplementary Tables S2 and S3). Further study in a logistic regression model with confounding factors and TR as a response variable – for each of the ten variables separately – limited the number of factors to be approved in the multivariable modelling to four factors (P <0.05) (Supplementary Tables S2, S4). These factors were gingivitis, stomatitis, periodontitis and dental calculus.

**Table 1.** Demographics and basic association test results (breed excluded) of cats with and without veterinarian-diagnosed tooth resorption (only P < 0.2 shown) in a subpopulation of 944 dentally examined and sedated cats from a feline health survey (data collected during 12/2012**–**2/2015). Number (n), percentage (%), CI = confidence interval, p-value: Fisher or Chi-square test. The same cat may have had several oral/dental diseases or viral infections. Bolded: selected for further analysis in a logistic regression model with confounding factors age, gender, and breed. The high proportion of missing values prevented the inclusion of vaccinations in further analyses.

| Variable | Diagnosed |  |  | Not diagnosed |  |  | P-value |
| --- | --- | --- | --- | --- | --- | --- | --- |
|  | N | % | 95% CI | N | % | 95% CI |  |
| **Age** | 201 |  |  | 730 |  |  | **<0.001** |
| < 7 yr | 51 | 25.4 | 19.9–31.8 | 295 | 40.4 | 36.9–44.0 |  |
| 7–10.99 yrs | 82 | 40.8 | 34.2–47.7 | 234 | 32.1 | 28.8–35.5 |  |
| ≥11 yrs | 68 | 33.8 | 27.7–40.6 | 201 | 27.5 | 24.4–30.9 |  |
| **Sex** | 199 |  |  | 736 |  |  | 1.000 |
| Female | 97 | 48.7 | 41.9–55.7 | 358 | 48.6 | 45.1–52.3 |  |
| Male | 102 | 51.3 | 44.4–58.1 | 378 | 51.4 | 47.8–55.0 |  |
| **Oral/dental disease** |  |  |  |  |  |  |  |
| Gingivitis | 74 | 36.6 | 30.3–43.5 | 194 | 26.2 | 23.1–29.4 | **0.005** |
| Stomatitis | 11 | 5.5 | 3.1–9.5 | 13 | 1.8 | 1.0–3.0 | **0.009** |
| Periodontitis | 22 | 10.9 | 7.3–15.9 | 35 | 4.7 | 3.4–6.5 | **0.002** |
| Dental calculus | 124 | 61.4 | 54.5–67.8 | 631 | 85.0 | 82.3–87.4 | **<0.001** |
|  | N | % | 95% CI | N | % | 95% CI |  |
| Malocclusion | 1 | 0.5 | 0.1–2.8 | 18 | 2.4 | 1.5–3.8 | 0.094 |
| **Diseases and disease categories** |  |  |  |  |  |  |  |
| Diseases of the nervous system | 5 | 2.5 | 1.1–5.7 | 8 | 1.1 | 0.6–2.1 | 0.166 |
| Parasites and protozoans | 18 | 8.9 | 5.7–13.6 | 44 | 5.9 | 4.5–7.9 | 0.148 |
| **Diet** |  |  |  |  |  |  |  |
| Cooked meat/fish | 59 | 29.2 | 23.4–35.8 | 177 | 23.9 | 20.9–27.1 | 0.120 |
| Constant availability of food | 108 | 54.3 | 47.3–61.1 | 446 | 61.0 | 57.4–64.5 | 0.088 |
| **Outdoor habits: cat outdoors** |  |  |  |  |  |  | 0.196 |
| All along | 39 | 19.4 | 14.5–25.4 | 137 | 19.1 | 16.4–22.2 |  |
| Every day | 40 | 19.9 | 15.0–26.0 | 149 | 20.8 | 18.0–23.9 |  |
| Weekly | 50 | 24.88 | 19.4–31.3 | 129 | 18.0 | 15.4–21.0 |  |
| Monthly | 28 | 13.93 | 9.8–19.4 | 101 | 14.1 | 11.7–16.8 |  |
| Less often than monthly/never | 44 | 21.89 | 16.7–28.1 | 201 | 28.0 | 24.9–31.4 |  |

### Random forest

We achieved the best accuracy (0.84) for random forest classification by setting the number of candidate variables to 13 and the minimum number of observations in a terminal node to nine. The six most important factors in classification cats to TR and non-TR cats from an accuracy perspective were breed (importance=45.5), dental calculus (16.9), gingivitis (9.5), age group (9.5), periodontitis (6.2) and constant availability of food (6.2). All other factors except constant availability of food had significant association (p < 0.05) with TR in the single variable logistic regression models with confounding factors.

**Figure 1.**
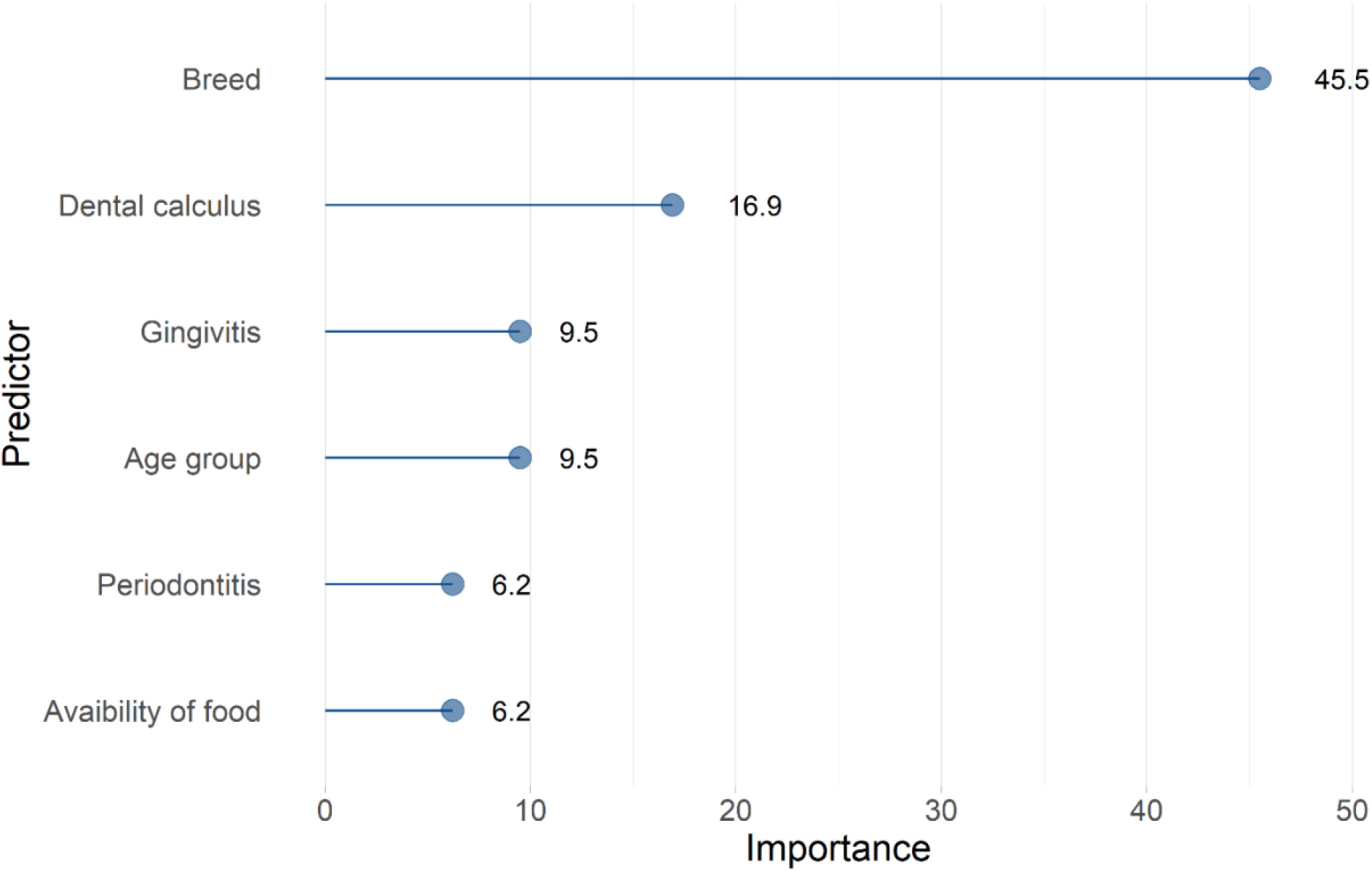
Results of random forest classification. Data: subpopulation of 944 dentally examined and sedated cats from a feline health survey (data collected during 12/2012–2/2015). Importance = importance of the predictor variables in classifying the data according to TR. The six most important variables presented.

### Multivariable logistic regression model and random forest classification

In addition to the confounding factors age, sex, and breed, five factors: gingivitis, stomatitis, periodontitis, dental calculus, and constant availability of food were qualified in the multivariable logistic regression analysis. Constant availability of food was included since it turned out to be one of the most important factors in RF classification, although it was not significant in factor-specific logistic regression analysis (p = 0.199) (Table 3). Based on the best model, independent risk factors for veterinarian-diagnosed TR were age and breed, which, in addition, were defined as confounding factors. Breeds with a significantly higher risk for tooth resorption than house cats were Cornish Rex,European and Ragdoll. Turkish Van and Devon Rex had no TR in the final model. Exotic-Persian group had significantly less TR than house cats.

Significant interactions with dental calculus were observed for gingivitis and periodontitis, and sex had interaction with constant availability of food. The dental disorder interactions demonstrated that gingivitis and periodontitis were risk factors for TR in the group of cats with dental calculus (Table 3). Female cats were significantly less prone to TR if they had food constantly available compared to those female cats who had feeding times. In male cats, we did not find that either feeding method affected TR susceptibility (Table 3).

The model achieved the best Pearson goodness of fit statistic, AIC and AUC values, and Cox and Snell index. Pearson’s goodness of fit statistics was 0.246. The AUC value for the ROC curve was 0.796 (95% CI 0.762 **–** 0.829), which makes the model’s predictive value moderate (Greiner et al., 2000). Cox and Snell goodness of fit index was 0.28. The goodness of fit and predictive value tests for the final model were mainly good or moderate.

When we compared the modelling in this study of 944 cats to similar modelling in the entire population of the feline health survey of 8115 cats (data not shown), results were parallel, except that stomatitis, different interactions and more breeds appeared to be risk factors in the entire feline health survey data.

**Table 2.** Multivariable logistic regression model of the risk factors for reportedly under sedation diagnosed tooth resorption in 944 Finnish cats. The data is a part of the cat health internet survey data collected 12/2012-2/2015. Only significant breeds are shown. B = logistic regression model coefficient, se = Standard Error, Wald = Wald test statistic, df = degrees of freedom, P = Wald’s P*-* value, OR = odds ratio, CI = confidence interval for OR, ref = reference group. n_cases_=196 and n_controls_=715.

| Variable | B | Se | Wald | P | OR | 95% CI | df |
| --- | --- | --- | --- | --- | --- | --- | --- |
| Constant | -0.04 | 0.35 | 0.01 | 0.92 |  |  | 1 |
| Sex: Male | -0.27 | 0.28 | 0.95 | 0.33 | 0.76 | 0.84–1.73 | 1 |
| Age: <7 | ref |  |  |  |  |  | 2 |
| 7-<11 | 0.81 | 0.23 | 12.61 | <0.001 | 2.25 | 1.44–3.51 | 1 |
| ≥11 | 0.92 | 0.25 | 13.72 | <0.001 | 2.51 | 1.54–4.07 | 1 |
| **Breed: House cat** | ref |  |  |  |  |  | 20 |
| Cornish Rex | 0.89 | 0.39 | 5.21 | 0.022 | 2.44 | 1.14–5.27 | 1 |
| European | 1.09 | 0.41 | 7.18 | 0.007 | 2.98 | 1.34–6.62 | 1 |
| Exotic-Persian | -1.27 | 0.59 | 4.61 | 0.032 | 0.28 | 0.09–0.90 | 1 |
| Ragdoll | 1.06 | 0.45 | 5.62 | 0.018 | 2.90 | 1.20–6.99 | 1 |
| **Interactions** |  |  |  |  |  |  |  |
| *Interaction gingivitis x dental calculus* | 1.98 | 0.43 | 21.44 | <.0001 |  |  | 1 |
| Gingivitis vs no when: |  |  |  |  |  |  |  |
| no dental calculus |  |  |  |  | 0.35 | 0.17–0.70 |  |
| dental calculus |  |  |  |  | 2.49 | 1.59–3.90 |  |
| *Interaction periodontitis x dental calculus* | 2.40 | 0.68 | 12.46 | <.0001 |  |  | 1 |
| Periodontitis vs no when: |  |  |  |  |  |  |  |
| no dental calculus |  |  |  |  | 0.33 | 0.11–0.98 |  |
| dental calculus |  |  |  |  | 3.70 | 1.62–8.45 |  |
| *Interaction food available constantly x sex* | 0.87 | 0.37 | 5.39 | 0.020 |  |  | 1 |
| Food available constantly vs no when: |  |  |  |  |  |  |  |
| Female |  |  |  |  | 0.44 | 0.26–0.75 |  |
| Male |  |  |  |  | 1.05 | 0.62–1.76 |  |

## Discussion

Our research on cat tooth resorption (TR) and its associated factors is based on a cross-sectional questionnaire survey of cat health that more than 8000 cat owners answered. We limited our study population to a subgroup of cats that had undergone a veterinary oral or dental examination under sedation to minimize potential false positives and negatives for TR in our study. We identified that dental calculus, gingivitis, and periodontitis as associated with TR and confirmed that the risk of TR increases with age. Breed-specificity was also observed, suggesting a genetic contribution to the aetiology of the disease.

The prevalence of TR in studies that have clinically evaluated the teeth is usually between 29–67% (Coles, 1990, Lund et al., 1998; Petterson and Mannerfelt, 2003; DeLaurier et al., 2009; Whyte et al., 2020; van Wessum, 1992; Lommer and Verstraete, 2000; Ingham et al., 2001). It is higher than the proportion of TR in our study population (21.4%). Our cross-sectional questionnaire study may expectedly result in low frequency because control cats may have had TR not found in sedation alone.

The examination methods and selection of the research population majorly impact the TR prevalence. The clinical studies of cats seeking dental treatment describe the prevalence of TR in the chosen study population rather than the prevalence in the entire cat population. If the target population consists of cats seeking dental treatment routinely or because of dental disease, the prevalence of TR is expected to be higher than in healthy cats. Most obvious lesions are found in the general examination without sedation, but examination with a dental probe under anaesthesia is required for most lesions. Only lesions in the cementoenamel junction or the crown area can be detected clinically. Dental radiographs increase the prevalence even higher when lesions in the root area are detected.

Varying results have been published considering the prevalence of TR in different breeds. Our results revealed that TR is highly associated with the breed – being purebred was not a risk factor, but the associations concerned certain breeds. We found Cornish Rex, European, and Ragdoll at higher risk for TR in the final multivariable model. In a recent study, Mestrinho et al. (2018) found a higher frequency of TR in Persian and Exotic cats. Our study, on the other hand, found less TR than average in these breeds. In addition, there was no TR in the Turkish van and Devon rex breeds. The breed predisposition might indicate a genetic component in the aetiology of the disease. Some breeds might be genetically predisposed to dental and oral diseases in general.

We found breed, age, dental calculus, gingivitis, periodontitis and constant availability of food to be associated with TR. These results were confirmed in both multivariable logistic regression modelling and random forest classification.

Our findings might suggest that dental calculus causing gingivitis and periodontitis increases the risk of TR. Gorrel’s (2015) theory suggests that tooth resorption consists of at least two aetiologically different diseases: inflammatory type 1 resorption and idiopathic type 2 resorption. Tooth resorption lesions often include inflammation of the adjacent gingiva (Reiter et al., 2005a; Pistor et al., 2023), making it difficult to assess causation between TR and gingivitis. Bacteria in dental plaque may initiate inflammatory resorption (Booij-Vrieling et al., 2010). Dental calculus causes gingivitis in cats, as reported by Thengchaisri et al. (2017). In addition, dental calculus is most often essential in development of periodontitis (Harrel, 2022). However, in our data there were cats in which the owner had reported periodontitis but no dental calculus. This probably resulted from the fact that the veterinarian diagnosed the disease but the owner answered the survey (see below limitations) or that the veterinarian had not reported the dental calculus in connection with diagnosis of periodontitis.

Periodontitis has been related to TR previously (Scarlett et al., 1999), and it has been linked especially with type 1 tooth resorption (DuPont and DeBowes, 2002). On the other hand, inflammation caused by tooth resorption has been suspected of causing periodontal lesions (Lemmons, 2013). A recent study by Whyte et al. (2020) found no relationship between TR and periodontitis. Although periodontitis would most likely be associated with inflammatory type 1 resorption, the type of TR was not determined in our study.

In our final multivariable model, stomatitis was not associated with TR. However, a low number of cats with stomatitis hindered us from studying this connection properly. DuPont and DeBowes (2002) found a connection between inflammatory type 1 resorption and stomatitis, but Reiter et al. (2005a) found no relationship. It was previously suspected that chronic stomatitis caused by the feline calicivirus impacts the development of TR (Reiter and Mendoza, 2002). However, Thomas et al. (2017) recently found that feline calicivirus (FVC) was associated with feline chronic gingivostomatitis but not with TR.

Our findings that the frequency of TR increases with age and sex not being associated with TR are in line with the previous studies (Coles, 1990; Harvey, 1993; Lund et al., 1998; Ingham et al., 2001; Pettersson and Mannerfelt, 2003; Reiter et al., 2005b; DeLaurier et al., 2009; Mestrinho et al., 2013; Pistor et al., 2023; Scarlett et al., 1999). In our study, sex was not associated with TR per se, but female cats were found to be less prone to TR when food was constantly available compared to feeding times. This was not seen in male cats. Scarlett et al. (1999) suspected indoor cats to have a higher risk of TR than outdoor cats, but Pettersson and Mannerfelt (2003) did not find a difference. We did not find a difference in the outdoor habits of the cats with or without TR. We could not reliably evaluate the effect of neutering and vaccinations due to uneven groups in neutering status and missing values in vaccinations. Considering other diseases, feline infectious peritonitis (FIP), cat flu (including herpes and calicivirus), leukaemia virus (FeLV), immunodeficiency virus (FIV) and feline panleukopenia virus were not associated with TR in our study.

Female cats that had food available constantly were less prone to TR compared to female cats that had feeding times. To the authors’ knowledge, the effect of food availability has not been studied before. This connection could be related to the consistency of the food. DuPont & DeBowes (2002) suspected dry food to cause mechanical trauma, leading to type 2 tooth resorption. These authors also suspected soft food, causing periodontitis, as a risk factor for inflammatory type 1 resorption. Scarlett et al. (1999) did not find a difference in prevalence between cats eating dry or soft food. In our questionnaire, we did not specifically ask if the cats ate dry or soft food. We could not find any explanation in our data to elucidate why constant availability of food in female cats would protect against TR. No connection was found between food availability and certain diet, outdoor habits or dental diseases. A further study would be needed to ensure whether food distribution habits have any effect on TR.

A limitation of our study is that we did not specifically ask about dental radiographs. It may have caused underestimations in our associations. In addition, although the veterinarian diagnosed the disease, the owner answered the survey. The owners might have understood, remembered, or interpreted the diagnoses in a way that could affect the results. Because the data were collected cross-sectionally, only the effect of permanent risk factors such as breed, age, and sex can be considered causal since they were permanently present before the disease. However, other factors found significant are potential risk or protective factors that should be verified with clinical trials or observational follow-up studies.

Interestingly, the results obtained in the subgroup limited and targeted to the study of TR correspond very well to the results of a similar study in the entire health and behavioural survey data. This confirms our understanding that the information from the cat health and behavioural survey gives a reliable overview of the health of the cats that participated in the survey.

## Conclusions

We have performed a cross-sectional questionnaire-based study of TR and identified several factors that were associated with TR. The risk of TR increases with age. Dental calculus, gingivitis, and periodontitis were associated with TR; however, it remains to be studied if these are true predisposing factors. These findings suggest that inflammatory changes caused by dental calculus might increase the risk of TR. Constant availability of food in female cats was found to be associated with TR, which also remains to be studied further. Finally, certain breeds appeared more predisposed to TR, suggesting a genetic contribution to the aetiology of the disease.

## Ethical statement

The data in this study were collected from 2012 to 2015 using an online feline health survey published by Vapalahti et al. (2016). The data were collected before the onset of the GDPR according to the Finnish legislation https://www.finlex.fi/fi/laki/ajantasa/1999/19990523. This survey study focused on investigating cats and not human participants or the cat owners; therefore, specific ethical approval was unnecessary. As for the study participants (cat owners), we collected only names and addresses. Owners were informed that the participation was voluntary and confidential and that the data was used only for scientific purposes. We received informed consent from all participants.

## Additional Information

## Acknowledgements

We thank the cat owners who participated in the original health survey. The HiLIFE and Jane and Aatos Erkko Foundation supported the study.

## Author contributions

HN, A-M.V. KV and HL conceptualized and designed the experiment. HN made the preliminary analysis and KV performed the multivariable analyses. KV and HN drafted the manuscript, which was edited and contributed by HL and A-M.V. All authors approved the final v**e**rsion.

## Supporting information

Data available: Supplementary Tables S1-S4 and S5.

## References

Ahola, M.K., Vapalahti, K., Lohi, H., 2017. Early weaning increases aggression and stereotypic behaviour in cats. Sci Rep. 7, 10412.

Booij-Vrieling, H.E., Ferbus, D., Tryfonidou, M.A., Riemers, F.M., Penning, L.C., Berdal, A., Everts, V., Hazewinkel, H.A.W., 2010. Increased vitamin D-driven signalling and expression of the vitamin D receptor, MSX2, and RANKL in tooth resorption in cats. Eur J Oral Sci. 118, 39–46.

Brown, L.D., Cat, T.T., DasGupta, A., 2001. Interval estimation for a proportion. Stat Sci. 16, 101–133.

Coles, S., 1990. The prevalence of buccal cervical root resorptions in Australian cats. J. Vet Dent. 7, 14–16.

DeLaurier, A., Boyde, A., Jackson, B., Horton, M., Price, J. 2009. Identifying early osteoclastic resorptive lesions in feline teeth: a model for understanding the origin of multiple idiopathic root resorption. J. Periodont Res. 44, 248–257.

Palmeira, I., Fonseca, M.J., Lafont-Lecuelle, C., Pageat, P., Cozzi, A., Asproni, P., Requicha, J.F., de Oliveira, J., 2022. Dental Pain in Cats: A Prospective 6-Month Study. J Vet Dent. Dec; 39(4):369–375.

DuPont, G., DeBowes, L., 2002. Comparison of periodontitis and root replacement in cat teeth with resorptive lesions. J. Vet Dent. 19, 71–75.

Farcas, N., Lommer, M., Kass, P., Verstraete, F., 2014. Dental radiographic findings in cats with chronic gingivostomatitis (2002-2012). JAVMA. 244, 339–345.

Gorrel, C., 2008. Small Animal Dentistry. Saunders Elsevier, 1st ed., China.

Gorrel, C., 2015. Tooth resorption in cats: patophysiology and treatment options. JFMS. 17, 37–43.

Gorrel, C., Larsson Å., 2002. Feline odontoclastic resorptive lesions: unveiling the early lesion. JSAP. 43, 482–488.

Greiner, M., Pfeiffer, D., Smith, R. D., 2000. Principles and practical application of the receiver-operating characteristic analysis for diagnostic tests. Prev Vet Med. 45, 23–41.

Harrel, S.K., Cobb, C.M., Sheldon, L.N., Rethman, M.P., Sottosanti, J.S., 2022. Calculus as a Risk Factor for Periodontal Disease: Narrative Review on Treatment Indications When the Response to Scaling and Root Planing Is Inadequate. Dent J (Basel). Oct 20;10(10):195. doi: 10.3390/dj10100195. PMID: 36286005; PMCID: PMC9600378.

Harvey, C., 1993. Feline dental resorptive lesions. Semin Vet Med Surg. 8, 187–196.

Ho, T. K., 1995. Random decision forests. In Proceedings of 3rd international conference on document analysis and recognition (Vol. 1, pp. 278–282).

Houe, H., Ersbøll, A. K., Toft, N., 2004. Introduction to Veterinary Epidemiology. Biofolia, Denmark.

Ingham, K., Gorrel, C., Blackburn, J., Farnsworth, W., 2001. Prevalence of odontoclastic resorptive lesions in a population of clinically healthy cats. JSA. 42, 439–443.

Kuhn, M., 2022. caret: Classification and Regression Training. R package version 6.0-91. https://CRAN.R-project.org/package=caret.

Lemmons, M., 2013. Clinical feline dental radiography. Vet Clin Small Anim. 43, 533–554.

Liaw, A., Wiener, M., 2002. Classification and Regression by randomForest. R News 2(3), 18–-22.

Lommer, M.J., Verstraete F.J.M., 2000. Prevalence of odontoclastic resorption lesions and periapical radiographic lucencies in cats: 265 cases (1995-1998). JAVMA. 217, 1866–1886.

Lund, E., Bohacek, L., Dahlke, J., King, V., Kramek, B., Logan, E., 1998. Prevalence and risk factors for odontoclastic resorptive lesions in cats. JAVMA. 212, 393–395.

Mestrinho, L., Runhau, J., Braganca, M., Niza, M., 2013. Risk assessment of feline tooth resorption: a portuguese clinical case control study. J. Vet Dent. 30, 78–83.

Mestrinho, L.A., Lauro, J.M., Gordo, I.S., Niza, M.M.R.E., Requicha, J.F., Force, J.G., Gawor, J. P., 2018. Oral and dental anomalies in purebred brachycephalic Persian and Exotic cats. JAVMA. 253, 66–72.

Okuda, A., Harvey, C., 1992. Etiopathogenesis of feline dental resorptive lesions. Vet Clin Small Anim. 22, 1385–1403.

Pett, M. A., 1997. Nonparametric Statistics for Health Care Research: Statistics for Small Samples and Unusual Distributions. Sage Publications, United States.

Pettersson, A., Mannerfelt, T., 2003. Prevalence of dental resorptive lesions in Swedish cats. J. Vet Dent. 20, 140–142.

Pistor, P., Janus, I., Janeczek, M., Dobrzyński, M., 2023. Feline Tooth Resorption: A Description of the Severity of the Disease in Regard to Animal’s Age, Sex, Breed and Clinical Presentation. Animals (Basel). Aug 3;13(15):2500

R Core Team, 2019. R: A language and environment for statistical computing. R Foundation for Statistical Computing, Vienna, Austria. URL https://www.R-project.org/.

Reiter, A., Mendoza, K., 2002. Feline odontoclastic resorptive lesions – an unsolved enigma in veterinary dentistry. Vet Clin Small Anim. 32, 791–837.

Reiter, A., Lewis, J., Okuda, A., 2005a. Update on the etiology of tooth resorption in domestic cats. Vet Clin Small Anim. 35, 913–942.

Reiter, A., Lyon, K., Nachreiner, R., Shofer, F., 2005b. Evaluation of calciotropic hormones in cats with odontoclastic resorptive lesions. AJVR. 66, 1446–1452.

Salonen, M.K., Vapalahti, K., Mäki-Tanila, A., Lohi, H., 2019. Breed differences of heritable behaviour traits in cats. Sci Rep. 9, 7949.

Scarlett, J.M., Saidla, J., Hess, J., 1999. Risk factors for odontoclastic resorptive lesions in cats. JAAHA. 35, 188–192.

Thengchaisri, N., Steiner, J.M., Suchodolski, J.S., Sattasathuchana, P., 2017. Association of gingivitis with dental calculus thickness or dental calculus coverage and subgingival bacteria in feline leukemia virus- and feline immunodeficiency virus-negative cats. Can J Vet Res. 81, 46–52.

Thomas, S., Lappin, D.F., Spears, J., Bennett, D., Nile, C., Riggio, M.P., 2017. Prevalence of feline calicivirus in cats with odontoclastic resorptive lesions and chronic gingivostomatitis. Res Vet Sci. 111, 124–126.

van Wessum, R., Harvey, C., Hennet, P., 1992. Feline dental resorptive lesions prevalence patterns. Vet Clin Small Anim. 22, 1405–1415.

Vapalahti, K., Virtala, A-M., Joensuu, T., Tiira, K., Tähtinen, J., Lohi, H., 2016. Health and behavioral survey of over 8000 Finnish cats. Frontiers Vet Sci. 3, 70.

Whyte, A., Lacasta, S., Whyte, J., Vicente Monteagudo, L., Tejedor, M.T., 2020. Tooth Resorption in Spanish Domestic Cats: Preliminary Data. Top Companion Anim Med. 38, 100369.

